# Genetic differentiation in *Anopheles stephensi* from India and Ethiopia: insights from mitochondrial genome analysis

**DOI:** 10.1101/2025.04.28.650555

**Authors:** Bhavna Gupta, Melveettil Kishor Sumitha, Mariapillai Kalimuthu, G NavaneethaPandiyan, Ramalingam Natarajan, Sampath Gopalakrishnan, Rajaiah Paramasivan, Manju Rahi, Ashwani Kumar

## Abstract

*Anopheles stephensi* is a primary malaria vector, particularly in urban areas of South Asia and the Middle East. Its recent spread into Africa has raised significant interest in understanding its genetic structure and evolutionary history. This study presents a comprehensive analysis of 98 mitochondrial genomes of *An. stephensi* from India, Pakistan, and Ethiopia, providing critical insights into the species’ genetic diversity patterns. High genetic diversity was observed among Indian samples, with two distinct genetic ancestries identified. While one ancestry was more prevalent in specific geographical regions, both lineages were found across all study sites, suggesting the co-existence of multiple lineages within these areas. Further analysis revealed significant genetic differentiation between Indian and Ethiopian populations, with limited overlap in shared mutations. Although some Ethiopian samples showed genetic relatedness to the SDA500 strain from Pakistan (used in genome sequencing by the Broad Institute), the majority of Ethiopian samples showed distinct genetic ancestry. The distinctiveness of Ethiopian populations, compared to other countries outside of Africa, was also reflected in the COI datasets analyzed in this study. However, further expanding the mitogenome dataset to include additional wild samples from regions such as Pakistan, Iran, Saudi Arabia, Sri Lanka and other African countries will provide a more comprehensive understanding of *An. stephensi*’s invasion history. Overall, the findings of this study highlight the significant genetic differentiation between Ethiopian and Indian *An. stephensi* populations, and existence of multiple lineages in Ethiopia indicating multiple independent introductions from different countries.

## Introduction

Globalization and increasing international trade have facilitated the movement of many species from their native ranges to new areas. While not all species can survive and establish themselves in these new environments, mosquitoes have shown remarkable resilience. The changing climate and rapid urbanization are making suitable breeding sites for their adaptation and survival. For example, *Aedes aegypti* and *Aedes albopictus*, both major vectors of diseases such as dengue and Zika, have expanded their geographical ranges dramatically in recent decades, often facilitated by human activities and urbanization [1,2]. A more recent and concerning addition to this list is *An. stephensi*, a primary urban vector of malaria in South Asia and the Middle East, which has recently been reported in several African countries rapidly expanding its range and posing new challenges for malaria elimination goals.

First detected in Djibouti in 2012, *An. stephensi* has since spread rapidly across the region [3]. In Ethiopia, it was initially found in Kebri Dehar in the Somali region [4], and by 2021, the species had expanded to multiple sites across the country. Subsequently, *An. stephensi* was reported in Sudan in 2019 [5,6], Nigeria in 2020 [7], Yemen in 2021 [8], Kenya in 2022 [9], Eritrea in 2022 [10] and Ghana in 2023 [11]. Unlike the primarily rural malaria vectors *An. gambiae* and *An. arabiensis* in sub-Saharan Africa [12,13], *An. stephensi* thrives in urban areas, breeding in artificial containers. This adaptation allows it to proliferate in rapidly urbanizing regions, representing a novel challenge for malaria control in sub-Saharan Africa, where urbanization is accelerating, and traditional vector control measures may prove less effective.

The recent spread of *An. stephensi* in Africa has sparked significant interest in understanding its genetic diversity and the sources of its introduction predicting its spread across international borders. Despite being an important malaria vector in South Asia, *An. stephensi* remains understudied genetically compared to other malaria vectors like *An. gambiae*. Although several studies have been published on African populations recently, genetic data on *An. stephensi* is limited, particularly from its native range such as India, Pakistan, Iran and Saudi Arabia. The studies on African populations have focused primarily on identifying *An. stephensi* and finding the possible source of origin. Global comparisons of COI sequences have shown matches with sequences from Pakistan and Sri Lanka [14], and some haplotypes from Sudan were found to be closely related to those from Saudi Arabia [15]. Although samples from various African countries, including Kenya, Djibouti, Sudan, Ethiopia, and Yemen, appeared genetically identical [16,17], this suggests the potential spread of *An. stephensi* within the continent, likely facilitated by extensive regional connectivity and trade. However, the lack of comprehensive data covering the entire geographic range of *An. stephensi* has complicated efforts to identify the precise origin of its invasion. Additionally, while the COI barcode region is a commonly used tool, it has limitations and may not be effective for detecting finer genetic variations. Recent genome-wide analysis of Ethiopian samples has revealed substantial genetic diversity, pointing to the establishment of a complex local population structure [14,17,18]. The identification of distinct populations in the southeastern regions, separate from those in other areas, suggests multiple independent introductions of *An. stephensi* into Ethiopia. This underscores the need for larger, more comprehensive datasets, which could provide a detailed understanding of the species’ genetic diversity and evolutionary history, especially if data from its entire distribution range are included in future studies.

In this context, we analysed a mitochondrial genome dataset from 98 samples, including those from India, Pakistan, Ethiopia, and laboratory colonies of Indian and Pakistan strains. This dataset represents a valuable resource for future genetic comparisons and offers critical insights into the genetic diversity of *An. stephensi*. Combining this data with global populations through an international research network, a more comprehensive picture of the species’ evolutionary history can be constructed. This collaborative effort will not only enhance our understanding of *An. stephensi*’s spread and invasion patterns but also help inform strategies to limit its further expansion.

## Materials and Methods

### Sample processing for genome sequencing

Five mosquito samples collected from Madurai and Trivandrum in India were analyzed in this study. Individual mosquito was used for DNA extraction using the DNeasy Blood and Tissue Kit (Qiagen, Cat# 69506), following the manufacturer’s protocol. The purity and concentration of the extracted genomic DNA were quantified using a Nanodrop Spectrophotometer (Thermo Scientific; 2000) and a Qubit dsDNA HS Assay Kit (Thermo Fisher Scientific; Cat# Q32854), respectively. Samples with optimal DNA yield and concentration (>2 ng/μL) were selected for further processing. Approximately 50ng of Qubit-quantified DNA was enzymatically fragmented using the FX Enzyme Mix provided in the QIASeq FX DNA kit Library Preparation protocol (Qiagen, Cat# 180475), following the manufacturer’s instructions.

The fragmented DNA underwent end-repair and A-tailing, followed by adapter ligation. Index-incorporated Illumina adapters were ligated to the fragments, generating sequencing libraries. The libraries were then subjected to six cycles of Indexing-PCR with the following thermal cycle parameters: Initial denaturation at 98°C for 2 minutes, followed by 6 cycles of denaturation at 98°C for 20 seconds, annealing at 60°C for 30 seconds, extension at 72°C for 30 seconds, and a final extension at 72°C for 1 minute. The amplified libraries were purified using Cambrian Bioworks CamSelect NGS (CBWD010), followed by quality control using the Qubit fluorometer (Thermo Fisher Scientific, MA, USA) and Agilent 4150 TapeStation for fragment size distribution analysis. Sequencing was performed by Genotypic Technologies (Bengaluru) on an Illumina NovaSeq 6000 platform using 150 bp paired-end chemistry.

### Mitogenome assembly and annotation

A total of approximately 42 to 67 million Illumina paired-end reads were generated from each mosquito sample. Raw reads were processed to remove adapters and low-quality reads using Trim Galore v.0.6.104 [19]. High-quality, adapter-clipped reads were mapped to the reference mitochondrial genome of *An. stephensi* (Accession No. GCF_013141755.1, 243.46 MB) using Bowtie 2 [20]. Mapped reads were assembled into the mitochondrial genome using NOVOPlasty [21]. The assembled mitochondrial genomes were annotated with MITOS [22] to identify the coding sequences for protein-coding genes and rRNAs. Sequence read archive (SRA) data for 90 samples were retrieved from the SRA database (IDs) and mitochondrial genomes were similarly assembled and annotated using the same workflow. Ethiopia, Pakistan, and laboratory colonies of Indian and Pakistan strains were included. The complete bioinformatics workflow, from DNA extraction to sequencing and analysis, is illustrated in Figure 1.

**Figure 1.**
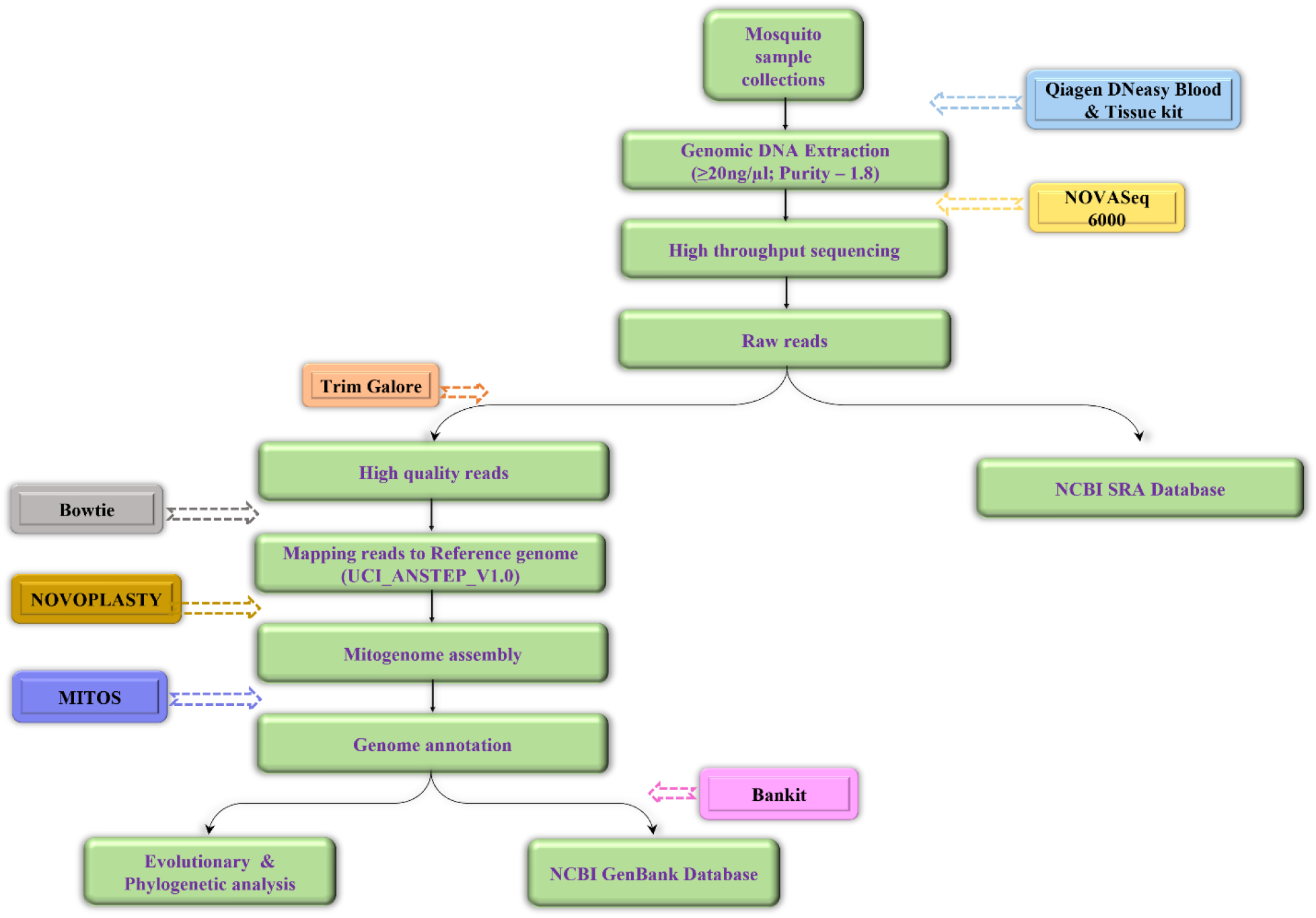
Bioinformatics workflow for mitogenome analysis of *Anopheles stephensi*, spanning from DNA isolation to genome sequencing, assembly, and downstream analysis

### Genetic analyses of mitochondrial data

Two separate datasets were analyzed, one using complete mitochondrial genomes and the other using the COI region, incorporating all available sequences from GenBank (Supplementary Tables S1, S2, S3). Sequences were aligned using MUSCLE in MEGA7 [23] and variants were identified. The number of polymorphic sites, the values of neutrality tests such as Tajima’s D, Fu & Li’s D were estimated using DnaSP [24]. Haplotype identification was performed using DnaSP. Minimum-spanning haplotype networks were constructed using NETWORK v10.0 (using median-joining network). Population structure analysis was performed with STRUCTURE software [25] using Markov Chain Monte Carlo (MCMC) methods, with burn-in set to 200,000 and 600,000 MCMC iterations, assuming correlated allele frequencies under the admixture model.

Phylogenetic analysis of the mitochondrial genomes was performed using MEGA7 with maximum-likelihood (ML) methods and bootstrap values from 500 replicates. The Tamura 3-parameter model with uniform rates was selected as the best-fit model of molecular evolution based on the Akaike Information Criterion (AIC). All protein-coding genes and rRNAs were aligned separately using MUSCLE, and the resulting alignments were analyzed for genetic variation, haplotypes, and haplotype diversity. All the analysis including haplotype network, phylogenetic analysis and STRUCTURE included both polymorphic sites (SNPs and indels) as variations.

## Results

### *An. stephensi* mitogenome sequences

Whole genome resequencing was performed on five *An. stephensi* samples from India, generating 46 to 67 million raw reads. Following quality control, adapter removal, and trimming, 40 to 64 million high-quality reads were obtained, resulting in 93 to 97% of the original reads being retained. These high-quality reads were then mapped to the reference mitochondrial genome of *An. stephensi* (NC_028223.1), with an average read depth of 30X. No ambiguous bases or heteroplasmic sites were detected in any of the samples.

Additionally, mitogenome assemblies were performed on 90 datasets obtained from the SRA database (SRA IDs are given in Supplementary Table S1), which contributed to generating additional *An. stephensi* mitogenomes. These assemblies were submitted to GenBank under accession number (submitted in GenBank). All assembled mitogenomes were circular, with an average length of approximately 15,380 bp, which is comparable to the reference mitogenome (NC_028223.1, 15,387 bp). Three reference mitochondrial genomes (Supplementary Table S1) were also included in the dataset resulting in 98 mitogenomes for final analysis. The mitochondrial genome of *An. stephensi* consisted of 37 genes, including 13 protein-coding genes (PCGs), 2 rRNAs, and 22 tRNAs (Table 1).

**Table 1.** Mitochondrial genes, their length, and the number of singletons, and nonsynonymous mutations found in the 98 mitochondrial genomes of *Anopheles stephensi*.

|  | **Length** | **SNPs** | **Singletons** | **Non-synonymous/singletons** |
| --- | --- | --- | --- | --- |
| **12S** | 687 | 5 | 5 | NA |
| **16S** | 1334 | 8 | 7 | NA |
| **ND1** | 1026 | 10 | 4 | 4/2 |
| **CYTB** | 1135 | 6 | 2 | 2/1 |
| **ND6** | 522 | 2 | 1 | 1/1 |
| **ND4L** | 297 | 4 | 2 | 2/1 |
| **ND4** | 1344 | 5 | 3 | 2/1 |
| **ND3** | 329 | 3 | 0 | 3/0 |
| **COX3** | 788 | 5 | 4 | 3/3 |
| **ATP6** | 681 | 3 | 1 | 0 |
| **ATP8** | 162 | 0 | 0 | 0 |
| **COX2** | 685 | 5 | 0 | 2/0 |
| **COX1** | 1537 | 12 | 5 | 2/0 |
| **ND2** | 1026 | 2 | 0 | 1/0 |
| **ND5** | 1743 | 16 | 9 | 10/6 |

Among the 98 samples analyzed, the ND5 gene exhibited the highest number of single nucleotide polymorphisms (SNPs) and non-synonymous mutations, followed by the COI gene (Table 1). Both ND5 and COI contained the maximum number of parsimony-informative sites. ATP8 showed no SNPs, while the SNPs found in the 12S and 18S rRNA genes were singletons. Out of the total of 108 SNPs identified, 86 were located in the protein-coding genes, 18 in the AT-rich region, and the remaining four SNPs were found in other regions of the genome.

### Genetic diversity among Indian *An. stephensi* samples

Five *An. stephensi* mitogenomes sequenced in this study were analysed along with 40 mitogenomes generated from publicly available genome sequencing data [26], which included 20 wild-caught mosquitoes and 20 from laboratory colonies. In total, 42 polymorphic sites (SNPs and Indels) were identified across 45 samples, resulting in 19 distinct haplotypes. STRUCTURE analysis revealed two genetic ancestries using Evanno’s method of finding the best k based on delta K values (Figure 2a). Seven wild-caught samples from Mangalore and two each from Madurai and Trivandrum clustered together, while seven wild-caught samples from Bengaluru formed a second cluster. Laboratory colony samples from Bengaluru also clustered with the Bengaluru wild samples. In contrast, laboratory samples from Chennai, Delhi, and Mangalore exhibited mixed grouping.

**Figure 2.**
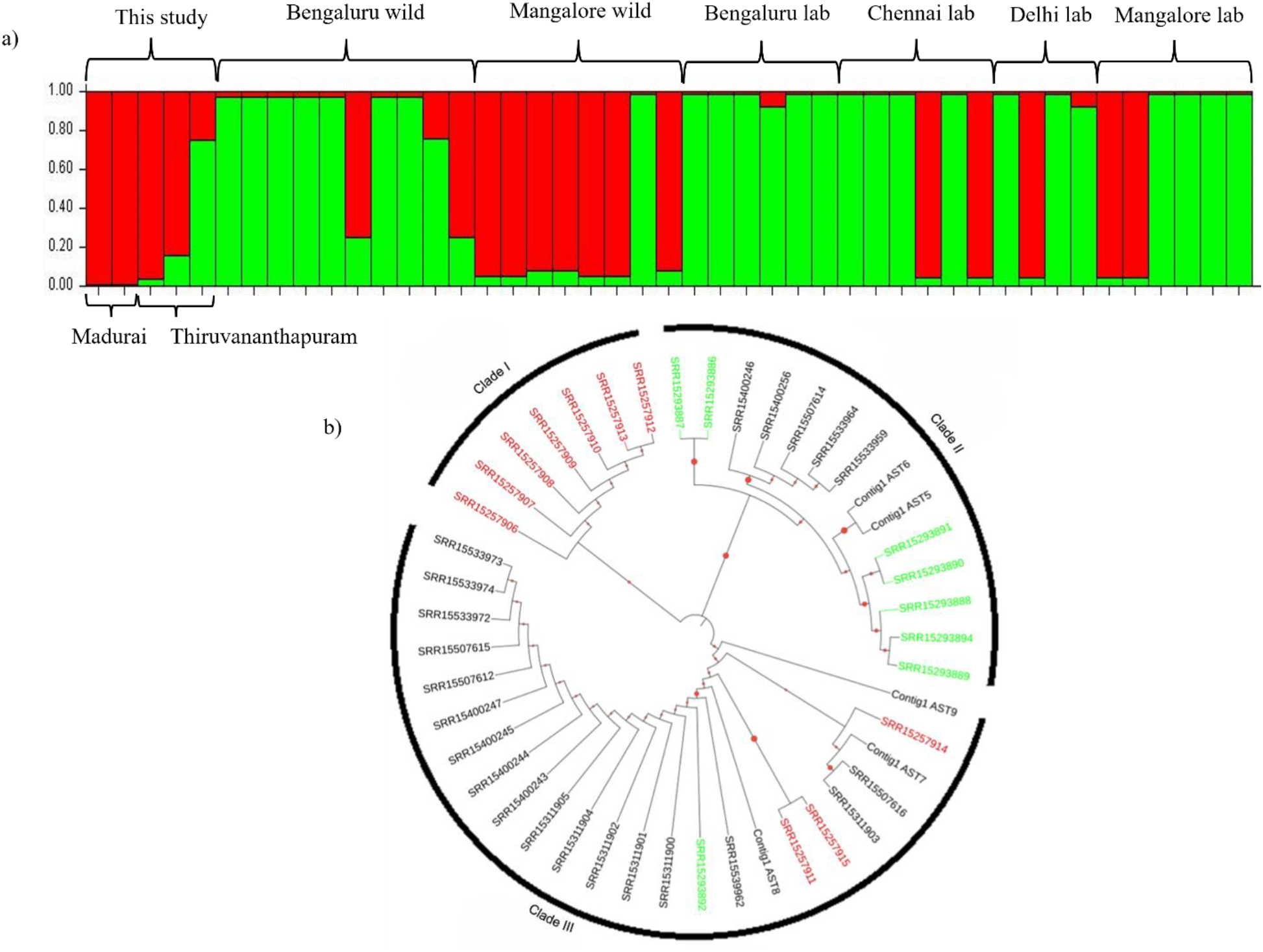
Genetic clustering analysis of Indian *An. stephensi* mitogenomes. a) STRUCTURE analysis indicating k=2 as the optimal number of genetic clusters. b) Maximum likelihood tree displaying the clustering of the *An. stephensi* mitogenomes based on genetic relationships (IDs in green represent samples from Bengaluru and red represents samples from Mangalore).

A similar pattern was observed in the phylogenetic tree, which showed three major clades (Figure 2b). While the majority of Bengaluru wild samples were found in Cluster 1, Mangalore and Madurai wild samples were predominantly grouped in Cluster II along with a few laboratory samples from Chennai, Delhi and Mangalore.

### Genetic diversity among Indian and Ethiopian samples

A total of 26 *An. stephensi* mitogenomes from Ethiopian samples were generated (one sample having several heteroplasmic sites was excluded from the analysis) [27], revealing 23 polymorphic sites resulting in 5 haplotypes. When combining the Ethiopian samples with Indian samples (71 mitogenomes in total), 57 polymorphic sites were identified, resulting in 29 haplotypes. Only eight mutations were shared between the Indian and Ethiopian samples.

Structure analysis using Evanno’s method identified k=2 as the optimal number of clusters (Figure 3a). However, considering the two clusters observed among Indian samples, k=3 (Figure 3b) provided additional insights. The Ethiopian samples were found to form a distinct group, with only three Ethiopian samples sharing genetic ancestry with Indian samples. A similar geographic division among mosquito samples was observed in the haplotype network (Figure 3c). Ethiopian haplotypes were clustered together, while Indian haplotypes showed extensive diversity. The presence of several median vectors within the Indian haplotypes represents potential ancestral or intermediate haplotypes that connect distinct genetic lineages, indicating a high level of genetic diversity that was not fully sampled in this study. Tajima’s D, Fu & Li’s D, and F values among Indian samples were -1.14, -1.39, and -1.55, respectively. In contrast, Tajima’s D among Ethiopian samples was -0.37, with Fu & Li’s D and F values of 0.88 and 0.60, respectively. Although all these values were statistically non-significant, the negative values observed in the Indian samples suggest a pattern consistent with population expansion and high genetic diversity.

**Figure 3.**
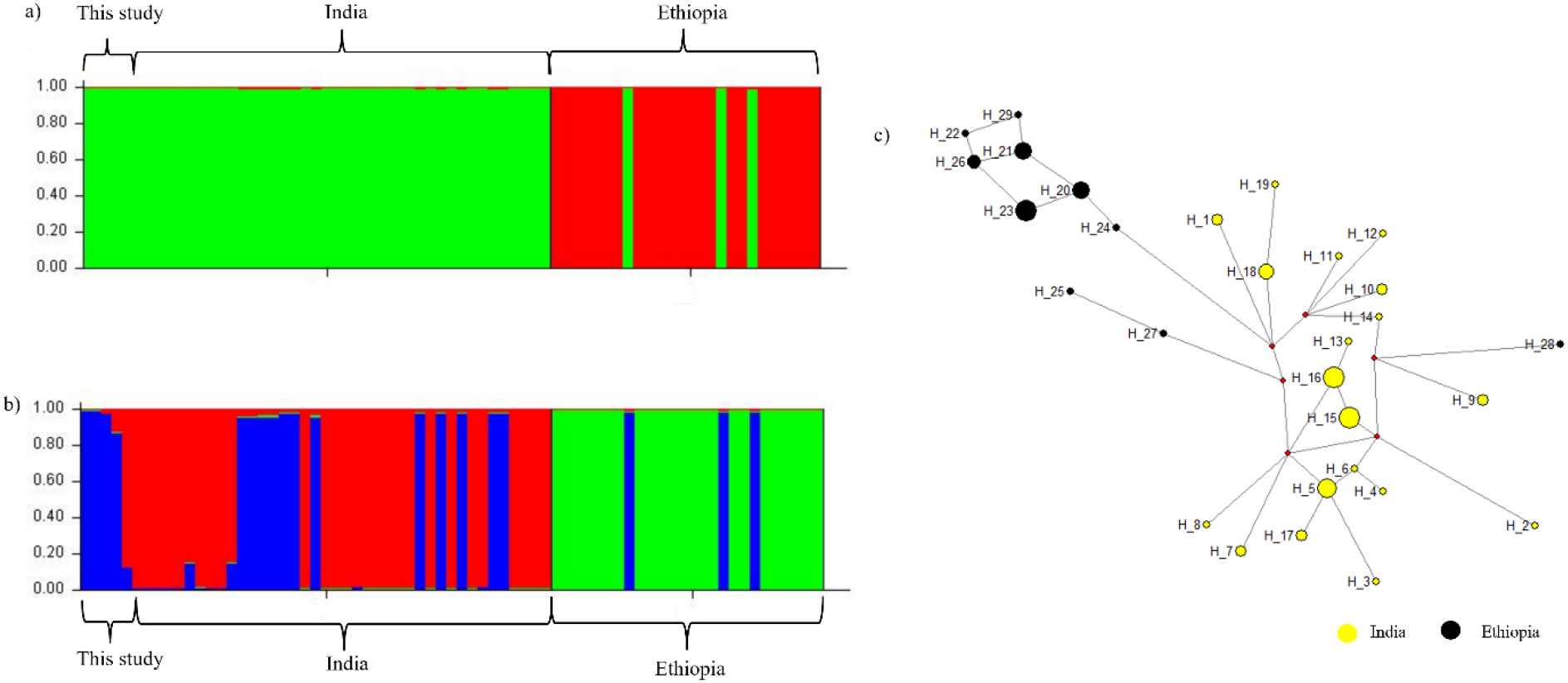
Genetic clustering analysis of Indian and Ethiopian *An. stephensi* mitogenomes. a) STRUCTURE analysis showing k=2 as the best model for genetic clustering. b) STRUCTURE analysis for k=3, revealing further genetic differentiation. c) Haplotype network constructed from 29 haplotypes, illustrating the relationships among different genetic variants.

Since the Ethiopian samples exhibited distinct genetic ancestry compared to the Indian samples, we expanded our dataset by incorporating 27 additional mitochondrial genomes from whole-genome sequencing (WGS) data available in the SRA database. This included 17 genomes from (laboratory colonies), 5 from Portugal (laboratory colonies), 2 from UK laboratory colonies, and 3 reference genomes. STRUCTURE analysis (k = 3) revealed that the Ethiopian samples clustered with the SDA500 strain from the Portuguese and UK laboratory colonies (Figure 4a). However, the SDA500 strain used for genome sequencing by the Broad Institute was found to be distinct and clustered with the Indian samples. Given the presence of several heteroplasmic sites in the Pakistani laboratory colony samples, we excluded 16 sites with heteroplasmy and trimmed the tail end that resulted into 87 variants (SNPs and indels). Re-running the STRUCTURE analysis on this filtered dataset revealed similar pattern showing the Ethiopian samples were genetically distinct from both the Indian and Pakistani samples, while showing closer genetic relatedness to the Portuguese and UK laboratory colonies, as observed in the full dataset (Figure 4b).

**Figure 4.**
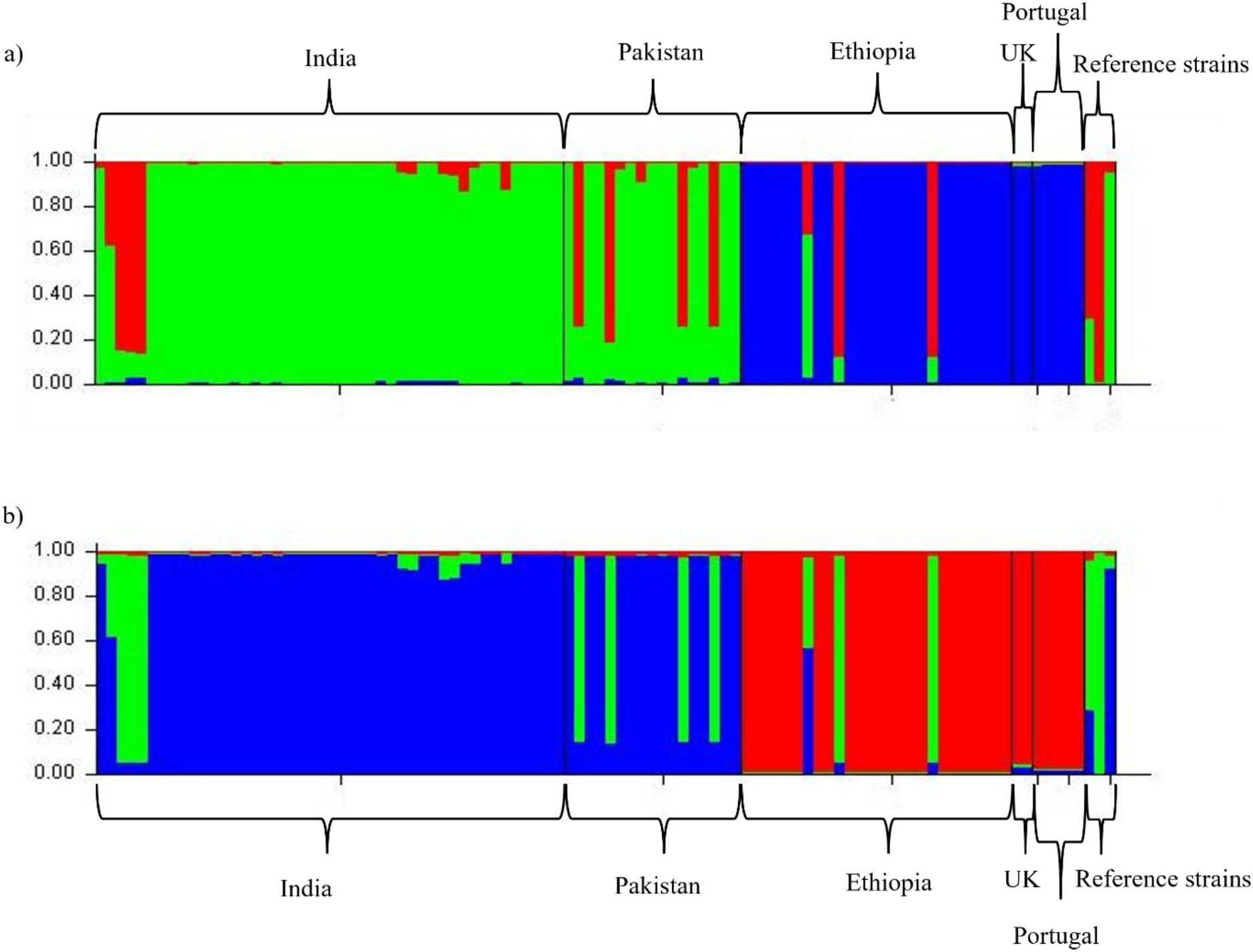
Genetic clustering analysis of *Anopheles stephensi* mitogenomes from global samples using STRUCTURE with k=3. a) Full dataset containing all variants. b) Filtered dataset after removing 13 heteroplasmic sites.

### Analysis of COI from global samples

To gain a global perspective on the genetic diversity of *An. stephensi*, we combined our dataset of 98 samples with publicly available COI sequences, resulting in a total of 533 samples of 237 bp (Supplementary Table 2). This combined dataset provided a broader view of the genetic variation within this mosquito species. The dataset included samples from South Asia (India, Pakistan, Sri Lanka), Africa (Kenya, Ethiopia, Sudan, Djibouti, Yemen, South Africa), and the Arabian Peninsula (Iran, Saudi Arabia, UAE). It is important to note that a significant portion of the samples (40%) in this dataset originated from Sudan. Additionally, we analyzed a longer stretch of the COI gene (435 bp) among 268 samples, which included samples from India, Pakistan, Sri Lanka, Ethiopia, Sudan, and Saudi Arabia (Supplementary Table 3). These COI datasets covered a wider geographical range compared to the full mitochondrial genome data.

In COI-smaller fragment-based dataset (hereafter called COI-I), a total of 15 single nucleotide polymorphisms (SNPs) and 19 haplotypes were identified, while longer fragment dataset (hereafter called COI-II) revealed 14 SNPs and 21 haplotypes. STRUCTURE analysis of the COI-1 dataset revealed three distinct genetic clusters (k=3), while the analysis of COI-II data (k=4) identified four clusters (Figure 5a, 5b). However, both the datasets showed similar pattern of genetic clustering among *An. stephensi* populations from different countries. For COI-I, majority of the samples exhibited two primary genetic ancestries in equal proportion (Figure 5a), with a third ancestry found only in samples from Sudan and Saudi Arabia. Some samples showed a higher proportion of one ancestry compared to the other; for example, a few samples from Pakistan, Ethiopia, and Sudan. Overall, most samples, regardless of their geographical origin, shared similar genetic ancestries, but the samples from Saudi Arabia and Sudan were genetically distinct. A separate cluster contained samples from Ethiopia, which exhibited genetic similarity to those from Pakistan and Sudan. In the COI-II dataset, South Asian samples shared common genetic ancestries, with the exception of a few samples from Pakistan that clustered more closely with African samples (Fig. 5b). Some samples from Ethiopia and Kenya were genetically closer to Asian samples, while the majority of Ethiopian and Kenyan samples were distinct, showing similarities only to two samples each from Pakistan, Sudan and the SDA500 strain from a laboratory colony in USA. Similar to the COI-I dataset, the samples from Saudi Arabia exhibited a unique genetic ancestry, further distinguishing them from other populations.

**Figure 5.**
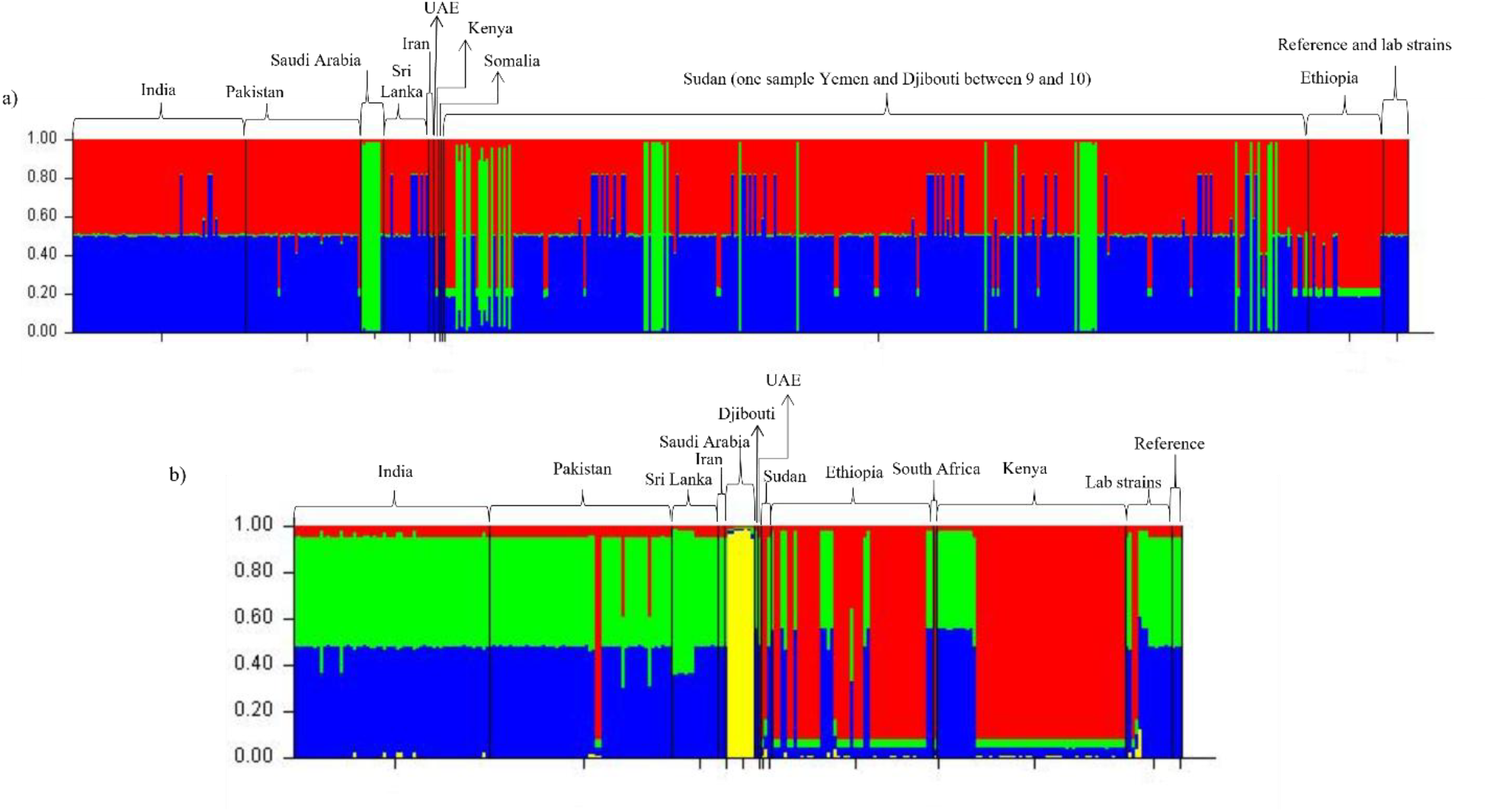
Genetic clustering analysis of *Anopheles stephensi* COI datasets from global samples using STRUCTURE. a) For COI-I samples with k=3. b) For COI-II samples with k=4.

## Discussion

One of the important contributions of this study is the generation of valuable genetic resource that will support future research into the evolutionary dynamics of *An. stephensi*. Although *An. stephensi* has been extensively studied in relation to its role in malaria transmission [28–30], its genetic diversity globally remains poorly understood in comparison to other mosquito vectors such as *An. gambiae* [31–33], *Ae. aegypti* [34–36], etc. In this study, we generated 95 mitochondrial genome assemblies from samples collected in India, Ethiopia, and other regions. This new dataset provides important insights into the molecular evolution and phylogeographic patterns of *An. stephensi*, shedding light on its genetic diversity and evolutionary history.

This is the first study to analyze the entire mitochondrial genome of *An. stephensi* populations across multiple countries. While several recent studies have focused on COI sequence comparisons across geographic ranges to identify the source of origin for newly invaded populations in Africa. Several interesting insights have bene gained from such studies. For example, comparisons of COI sequences globally revealed matches with sequences from Pakistan and Sri Lanka [14], and some haplotypes from Sudan were closely related to those from Saudi Arabia [15]. While samples from different African countries, Kenya, Djibouti, Sudan, Ethiopia, Yemen, etc. appeared genetically identical [16,17], suggesting a potential spread of the species within the continent due to the extensive connectivity and trade across the countries. However, the lack of comprehensive data from the entire geographic range of *An. stephensi* has made it difficult to pinpoint the exact source of invasion. Recent genome-wide variant analysis of Ethiopian samples [14,17,18] revealed considerable genetic diversity, indicating the establishment of a rich local population structure. The presence of distinct Southeastern populations, genetically separate from other regions, suggests the possibility of multiple independent introductions of *An. stephensi* into Ethiopia. This highlights the importance of larger datasets, which can provide a complete picture of the species’ genetic diversity and evolutionary history, especially if data from its entire range become available for future research.

In this study, we analyzed 98 mitochondrial genomes of *An. stephensi*, focusing on samples from India, Pakistan, and Ethiopia. Despite the limited geographic scope, our findings provided several key insights into the genetic structure of the species. Notably, high genetic diversity was observed among the Indian samples, which represent the species’ native range. Our STRUCTURE analysis revealed that all the Indian samples clustered into two main groups, but further examination of the haplotypes indicated substantial diversity within these groups. The diversity observed suggests a rich and complex genetic structure in the Indian populations of *An. stephensi*, consistent with previous studies that have identified high levels of genetic variation within the country [37–39]. Interestingly, while the samples from Bangalore and Madurai predominantly grouped into one genetic cluster, wild samples from Mangalore formed a distinct cluster. While both the lineages were observed in all the laboratory colonies and the field samples. This suggests the coexistence of different mitochondrial lineages of *An. stephensi* despite geographical distance between these locations. However, to fully capture the breadth of this diversity and understand the underlying evolutionary dynamics, it is crucial to expand the dataset to include more samples from other regions of India and beyond. Larger-scale studies that incorporate a broader geographical range would provide a more comprehensive understanding of the genetic landscape of *An. stephensi* and reveal whether these observed genetic lineages are geographically isolated or exhibit varying degrees of gene flow. This would also shed light on potential local adaptations and the role of ecological factors in shaping the species’ genetic diversity.

Interestingly, when Indian and Ethiopian *An. stephensi* samples were combined, the results underscored the complexity of the species’ genetic structure. Our analysis identified 43 SNPs and 29 distinct haplotypes across 71 mitogenomes, with only eight mutations shared between the two regions. Ethiopian samples displayed entirely distinct genetic ancestry, of which only three haplotypes showed genetic relatedness with Indian samples. This limited overlap suggests substantial genetic differentiation between Ethiopian and Indian populations. This geographic division aligns with the observations in the haplotype network, where Ethiopian samples appeared genetically distinct, with only a few sharing ancestries with Indian samples. The majority of Ethiopian-specific haplotypes exhibited only one or two SNP differences. In previous studies based on COI, Ethiopian haplotypes have been found to match with those from Pakistan, and Sri Lanka [4,27] similar patterns emerged in our combined analyses of laboratory samples from Pakistan and other regions. Although majority of Ethiopian samples remained distinct, three samples were found to be genetically closer to the Pakistani genetic cluster, suggesting some level of genetic connection between these regions. Intriguingly, Ethiopian samples showed a high degree of similarity to SDA500 laboratory colonies from Portugal and UK, while were distinct from the SDA500 strain samples used by Broad institute for whole genome sequencing. One possibility is that the genetic diversity in different laboratory colonies may have evolved over several generations under controlled conditions, leading to divergence from the original field populations. Additionally, the Pakistan SDA500 strain used by the Broad Institute may have originated from multiple field samples, which could explain the observed genetic divergence. Several heteroplasmic mutations were identified within the Pakistan samples, suggesting the presence of different mitochondrial haplotypes. Even after excluding heteroplasmic sites from the final analysis, the same genetic clustering pattern was observed, indicating a genetic relatedness between Ethiopian and laboratory colonies of SDA500 from Portugal and UK. However, the identification of new lineages in the Ethiopian samples is an important observation towards understanding the global distribution of *An. stephensi* and its potential for further spread across continents. However, the absence of comparable data from regions such as South Asia, the Arabian Peninsula, and other African countries limits our ability to construct a comprehensive picture of the species’ genetic diversity and evolutionary history. This highlights the need for future studies adding more diverse samples from these regions to this dataset to fully understand the global genetic landscape of *An. stephensi*.

Furthermore, to gain a global perspective, we revisited COI data from GenBank and combined our sequences with existing data, resulting in a dataset of 533 samples (237 bp). Three major clusters were identified: one containing samples from Pakistan, India, and Sri Lanka, the second containing samples from Saudi Arabia and Sudan, and the third consisting of samples from Ethiopia and Kenya. These regions also shared several common haplotypes. Analysis of a longer COI fragment revealed similar patterns, indicating that Ethiopia and Kenya had samples from two distinct groups: one that was genetically unique and the other that resembled samples from Pakistan. While samples from Sudan and Saudi Arabia formed distinct groups, they still showed some overlap with other clusters.

Although these observations are not entirely new, the inclusion of a larger sample size and the use of the complete mitochondrial genome offer additional insights into the genetic structure of *An. stephensi*. This comprehensive dataset serves as a valuable reference for future investigations into the origin and spread of *An. stephensi*. Expanding the sample collection to include more samples from its native range, including India, Pakistan, Iran, and Saudi Arabia, and comparing these with samples from African populations, would provide a clearer and broader picture of its invasion history. Such an analysis would greatly contribute to our understanding of the species’ dynamics, offering important information that could inform strategies to monitor and prevent its future spread.

This study provides valuable insights into the genetic diversity and evolutionary dynamics of *An. stephensi*, highlighting distinct mitochondrial lineages in Ethiopian populations and significant genetic differentiation from Indian samples. The analysis of mitochondrial genomes and COI sequences contributes a comprehensive genetic resource, revealing complex population structures and regional differentiation. However, the lack of data from underrepresented regions, such as the Arabian Peninsula and other parts of Africa, limits our understanding of its global genetic landscape. Expanding the dataset from these regions will offer a clearer picture of the species’ spread and evolution, informing future malaria control strategies and enhancing efforts to mitigate the spread of *An. stephensi* across continents.

## Acknowledgements

Authors would like to thank Indian Council of Medical research (ICMR), New Delhi for intramural support. MK Sumitha and G Navaneetha Pandiyan would like to thank Madurai Kamaraj University for supporting their research.

